# ClonalTracker: a tool to elucidate dissemination patterns between vancomycin-resistant *Enterococcus faecium* isolates

**DOI:** 10.1101/2024.01.05.574403

**Authors:** Victoria Pascal Andreu

**Affiliations:** Department of Medical Microbiology, Universitair Medisch Centrum Utrecht, Utrecht, The Netherlands

## Abstract

The global spread of vancomycin-resistant *Enterococcus faecium* (VRE), which commonly occurs in hospital environments, has become a major public health concern. To facilitate genomic surveillance and tracking the transmission of VRE, ClonalTracker was designed. This tool assesses the clonal relatedness between two VRE isolates given the respective assembled genomes by analyzing the *van* operon, the respective transposon type and the whole genome similarity. ClonalTracker has been validated using two previously analyzed publicly available datasets and I showcase its applicability on a yet unprocessed third dataset. While the method agrees with previously published results, it is able to provide more resolution at the clustering level even in the absence of plasmid information and using as reference the minimal version of the vancomycin resistance transposon. Within this third dataset composed of 323 *vanB* VRE isolates, ClonalTracker found that clonal expansion is the most common dissemination mode. All in all, this tool provides new bioinformatic means to uncover dissemination patterns and elucidate links between vancomycin-resistance isolates and can be broadly accessible via its webserver hosted at www.clonaltracker.nl (as of January 2024). The local version of this tool is also available at: https://github.com/victoriapascal/clonaltracker

## Introduction

*Enterococcus* species are ubiquitous in nature. They are especially common in the gastrointestinal tract of most mammals including humans, with two main players in terms of abundance: *Enterococcus faecium* and *Enterococcus faecalis*^1^. These species are known for their genome plasticity that may contribute to the disruption of their healthy association with the host given their ability to acquire new antimicrobial resistance and virulence genes^2^. In clinical settings, the intensive exposure of some patients to antibiotics resulted in the selection of antibiotic resistance in these *Enterococcus* species, most notably of vancomycin-resistance, particularly in *E. faecium*^3^. This has promoted the increase of life-threatening infections and their adaptation in hospital environments. Given this threat, the World Health Organization (WHO) has listed vancomycin-resistant *E. faecium* (VRE) as high-priority in their global compendium of important antibiotic-resistant bacteria^4^.

VREs have emerged as a major threat for public health especially in hospital settings^5^. This resistance is mediated by the acquisition of the *van* operon, which is composed of several genes including a ligase. There are different types of ligases giving name to different types of operons, that allow the substitution of peptidoglycan precursor D-Alanine-D-Alanine (D-Ala-D-Ala) to either D-Alanine-D-Lactate (VanA, VanB, and VanD ligases) or D-Alanine-D-Serine (VanC, VanE, and VanG ligases). The different *van* types have different gene contents and arrangements^6^ and provide different levels of resistance ranging from high (*van*A) to variable (*van*B)^7^. The resistance phenotype is the result of lower binding affinity of vancomycin for the modified cell-wall precursors^8^. It is known that this operon can be inserted in a transposon, which can integrate itself in different locations of the genome to later be transmitted clonally^9^. In addition, the *van*-encoding transposons can also be located on plasmids which, in the case of conjugative mobilizable plasmids, can be transferred horizontally between bacteria^10^.

Taking advantage of the growing number of genomes available online and with whole genome sequencing (WGS) being more implemented in healthcare-related laboratories, it is now possible to extract more information from outbreak-related isolates^11^ and can ultimately help contextualize and better interpret dissemination modes and patterns of antibiotic resistance. Different tools exist to extract different types of information from these genomes such as: core and accessory genes similarity between isolates^12^, predict antimicrobial resistance genes from WGS^13^ or detect single nucleotide polymorphisms (SNPs), insertions and deletions of the related mobile genetic elements^14^. However, there is no method that combines all these layers of information into one tool. Therefore, I have designed ClonalTracker, a Python3 framework to assess the pairwise relatedness of VRE isolates and provide insights regarding the *van* operon type, variation in the *van*-carrying transposon and overall genome similarity of VRE isolates. Moreover, to make the tool broadly accessible, ClonalTracker is also available as an online service at www.clonaltracker.nl. ClonalTracker pipeline has been validated and tested using two different datasets already analyzed and published^15,16^. The re-analysis of these two datasets showed that the predictions agree with the reported ones but is able to further classify isolates into more fine-grained outbreak clusters. The analysis of a third dataset composed of 323 *vanB* isolates showcases an application of how to use ClonalTracker and how to use its results for further analysis. These results provided insights on the spread of the *vanB* resistance genes and suggest that these are mainly clonally disseminated, with some indications of horizontal dissemination. Moreover, using some of the intermediate results I got further insights into the diversity and architecture of the *vanB*-harboring transposon.

### ClonalTracker pipeline

ClonalTacker is a Python3 tool that uses two assembled VRE genomes in FASTA format as input (https://github.com/victoriapascal/clonaltracker). It combines different tools to assess (1) the *van* type of each isolate, (2) their respective transposon type and (3) the whole genome similarity of both genomes, including the contextualization of the two genomes within a broader set of VRE genomes (see the overall workflow in Figure 1). First, it uses PopPUNK^12^, specifically the query assignment option, to fit the input genomes to a database composed of 656 *vanA* and *vanB* VRE genomes isolated from different Dutch hospitals between 1995 and 2015 (ENA accession number PRJEB28495). Next, the *van* type of both isolates is assessed using blastn^17^ paired with a database of 7 different types of D-Ala-(R) ligases including a representative of the *vanA, vanB, vanC, vanD, vanG, vanL* and *vanN* operons. If the ligase is detected and the two genomes have the same *van* type, the transposon typing is performed. As the tool has been designed to help survey hospital-related outbreaks, only *vanA* or *vanB* isolates (the most frequent types) can be fully analyzed. In case of other types, including genomes with more than one *van* operon, only the *van* typing step and PopPUNK analysis will be performed. The transposon typing step starts by gathering the contigs which align to either of the following reference transposon sequences: Tn1546 (accession M97297, 10851 bp) and Tn1549 (accession AY655721.2, 9841 bp), for *vanA* and *vanB* respectively using blastn. These transposon-like sequences include the *van* operon, the integrate and excisase. If the transposon is split into multiple contigs, RagTag^18^ is used to scaffold the transposon again using the corresponding reference sequence. Later, the start and end coordinates of the transposon are determined and used to trim the transposon sequence accordingly. To assess the IS insertions, ClonalTracker uses ISEScan^19^, followed by Bakta^20^ to annotate and generate the Genbank files used as input for visualization with clinker^21^ (version v0.0.24). The latter also performs all the pairwise protein sequence alignments between transposon’s’ coding genes, which are used also as a metric for transposon identity. Finally, to assess the transposon’s’ synteny and completeness, one of the two isolates’ transposons is randomly chosen and used to create a blast database with makeblastdb^22^. The other transposon sequence is used as a query to run blastn again. Lastly, for whole genome comparison the k-mer based approach Mash^23^ is used, sketching 8000 k-mers of size 32. To determine clonally-related isolates the Mash distances of the *van*A isolates were compared to the hierBAPS1-3^24^ sequence clusters (SC) and (Multi-Locus Sequence Typing)^25^ (https://github.com/tseemann/mlst) groups. Isolates sharing >95% of the Mash k-mers mostly belonged to the same ST/SC (134/144) and thus, are considered clonally-related isolates, while isolates sharing less than 90% of the k-mers but still sharing the same transposon are deemed candidates of horizontal transmission as they differed more at the ST/SC clustering level. Isolates sharing between 90-95% k-mers are dubious cases as the lack of similarity can be due to the randomness selection of k-mers or because they are truly more distantly related thus, they are classified as “unknown”. Although ClonalTracker uses two assembled genomes as input, the command line version can also be run within a *for* loop to perform all the pairwise comparisons.

**Fig 1.**
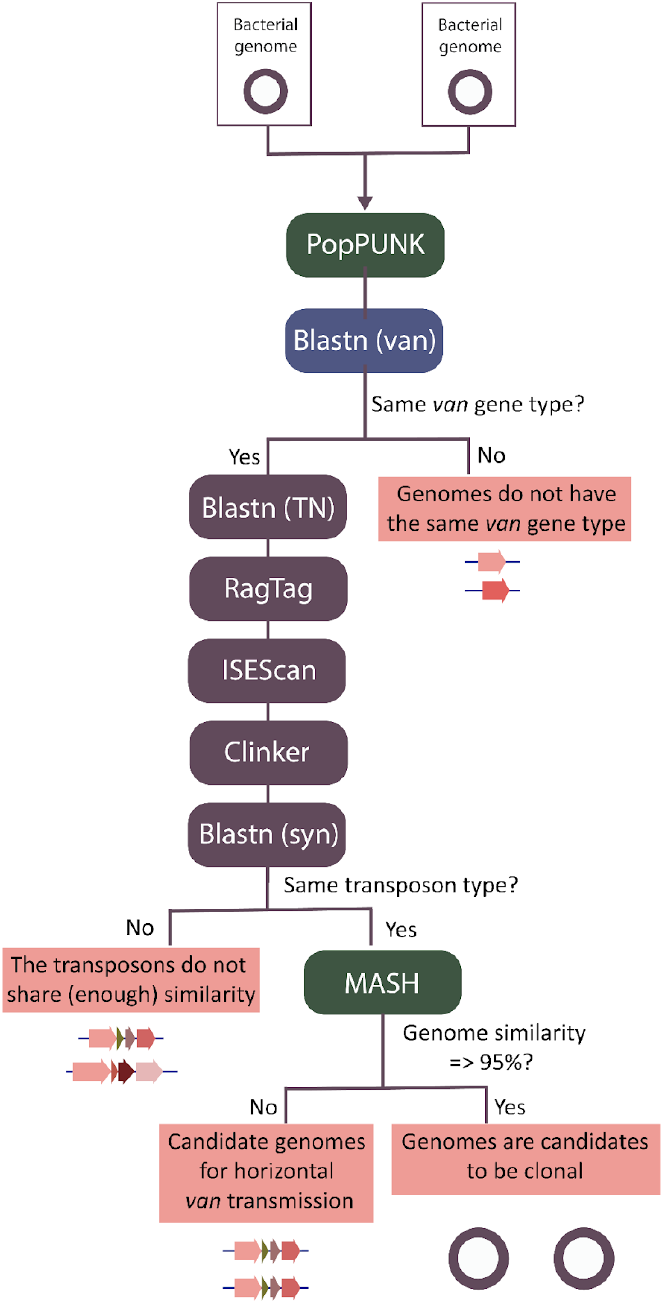
ClonalTracker workflow. The tool consists of three main steps: (1) *van* typing (blue box), (2) transposon typing (maroon boxes) and (3) whole-genome sequence similarity and contextualization (green boxes).

### Validation of ClonalTracker

ClonalTracker was validated using two publicly available datasets: 333 *van*A-VRE isolates from 31 different Dutch hospitals sequenced between 1995-2015^15^ (see Table S1) and 39 *vanB*-VRE isolates collected during an outbreak in 2014 in the University Medical Center Groningen (UMCG, The Netherlands)^16^ (see Table S2). The two datasets were downloaded from the European Nucleotide Archive (ENA) using the accession number PRJEB28495 and PRJEB25590, respectively. The raw reads were assembled using bactofidia (version 1.1, https://gitlab.com/aschuerch/bactofidia). To discern between clonal and Horizontal Gene transmission (HGT), I compared the Mash distances of each *vanA* pair of isolates with the MLST and hierBAPS1-3 clustering. This allowed to set up internal ClonalTracker thresholds to classify clonally-related and HGT-related isolates (see Methods: *ClonalTracker pipeline* for more details). To quantify the congruence between ClonalTracker’s classification method and MLST and hierBAPS the adjusted Wallace test^26^ was run. Finally, ClonalTracker’s results were visualized with Cytoscape^27^ and compared with the conclusions drawn from the initial analysis by Arredondo-Alonso et al^28^ and Zhou *et al*.,^16^, for the *vanA*-VRE and *vanB*-VRE dataset respectively.

### ClonalTracker webserver

The ClonalTracker webserver is available at www.clonaltracker.nl. The webserver has been designed using the Python3 Flask web framework (https://flask.palletsprojects.com) for server-side logic. The server allows any user to upload their FASTA files and automatically run the whole pipeline. Each run is identified with a unique job ID, and if an email address is provided in the submission step, the user will be notified with an email about the status of their submission. Otherwise, the job ID can be used to retrieve the results online. The user is able to visualize the results on the browser and also to download all the intermediate and final results. The webserver can nowadays process 6 submissions in parallel, and, depending on the workload, the job can be directly processed in about 10-15 minutes (from submission to completion). Otherwise, it will be queued until resources are available again.

## Results

### The ClonalTracker webserver facilitates VRE genome comparison

The ClonalTacker landing page (see Figure 2a) allows the user to submit a job providing two FASTA files as a starting point. Moreover, the webserver provides other functionalities such as retrieving the results given a job ID (“Results for existing job” tab) but also more information to better understand the ClonalTracker workflow (“Tutorial”) and a guide to fully understand all the results included in the interactive report generated by ClonalTracker (“Example output” tab, see Figure 2b). The results are fully interactive and downloadable.

**Fig 2.**
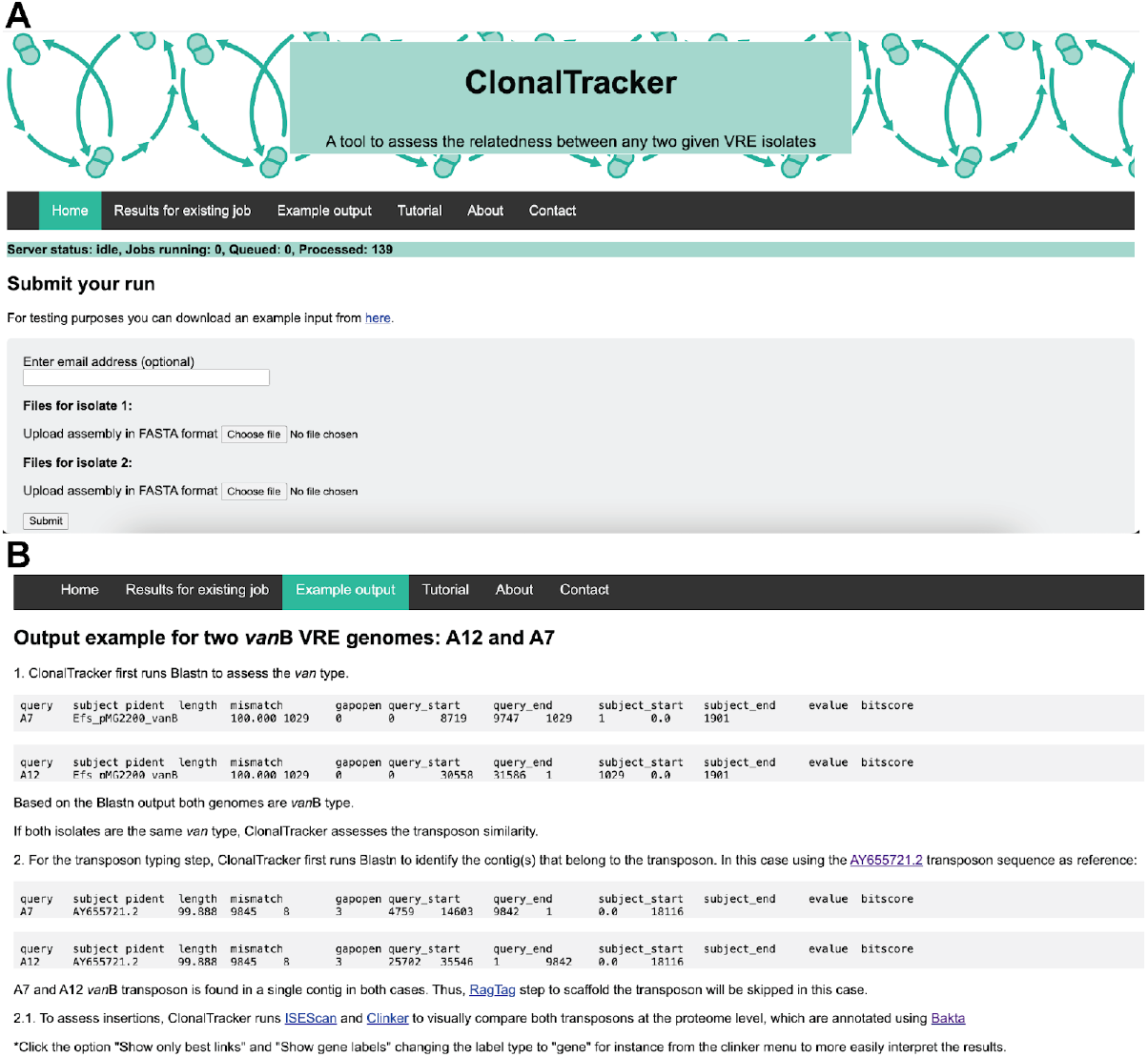
ClonalTracker webserver. a) ClonalTracker’s landing page to submit a new run and visualize the server workload. b) “Example output” tab provides information on the different results included in the final interactive report to help interpret the results and better understand them.

### Evaluation of the congruence between clustering methods

Two isolates are classified as clonally related when both genomes had the same *van* gene, transposon type and shared more than 95% of the Mash k-mers. To quantify the congruence between the MLST, hierBAPS1-3 and ClonalTracker clustering methods, the resulting clusters on *vanA* isolates from ClonalTracker were compared to those previously reported^28^ by MLST and hierBAPS1-3. First, the number of groups vary largely between clustering methods: 137 with ClonalTracker, 11 with MLST, 5 with hierBAPS1, 11 with hierBAPS2 and 18 with hierBAPS3. When comparing each clustering method to ClonalTracker’s clonal clusters, the adjusted Wallace coefficients were very low (see all results in Table S3). The highest value (0.133) was obtained for the hierBAPS3:ClonalTracker comparison (confidence interval of 0.09-0.177) and the lowest (0.046) for the hierBAPS1:ClonalTracker (confidence interval of 0.022-0.07). However, when ClonalTracker classification is compared to MLST and hierBAPS1-3 the values range between 1 (confidence interval of 1-1) for the ClonalTracker:MLST and ClonalTracker:hierBAPS1-2 comparisons and 0.960 (confidence interval of 0.905-1) for the ClonalTracker:hierBAPS3 comparison. These results suggest that while all methods agree in their classification despite finding significant differences in the granularity of their clustering, the direction in which clustering methods are compared does matter.

### ClonalTracker has more resolution at the clustering level

Two VRE datasets composed of (1) 333 *vanA* isolates and (2) 39 *vanB* isolates previously analyzed by Arredondo-Alonso et al^28^ and Zhou et al^16^ respectively were used to compare to ClonalTracker’s results. From the initial *vanA*-VRE dataset, five isolates were discarded for either not finding a *vanA* operon (n=3), having both a *vanA* and *vanB* (n=1, as the tool is not able to handle genomes with more than one *van* resistance gene) or being wrongly annotated as *vanA* (n=1) (see Table S1), resulting in a total of 328 curated isolates. For the *vanB*-VRE dataset, the whole population (n=39) was considered.

In the *vanA*-VRE dataset, ClonalTracker found that 164 isolates shared 35 different *vanA* transposons, while 164 isolates had an unique *vanA* transposon type, also known as singletons (see Figure 3 generated using Table S4). The network shows 164 nodes (isolates) connected by 1288 edges (links between isolates). From all the links, 854 represent clonal relationships between 144 isolates (see black edges in Figure 3), while 160 edges link 90 isolates with the same transposon but with a different genomic background (red edges). Compared to the already published results, ClonalTracker links more isolates compared to the original publication (49,1% compared to 41%). Both methods agree that within this dataset isolates are more often clonally-related than horizontally-related. However, Arredondo-Alonso et al^28^ also assessed the plasmid type often involved in the dissemination of the *vanA* resistance. To assess the impact of taking into account the plasmid type to further classify isolates, the classification of 225 isolates provided by the original publication were compared to the ones generated by ClonalTracker (see Table S5). The authors took into account the genomic background using hierBAPS (SC), transposon type retrieved by TETyper (TET) and plasmid type using Mash (PT), classification which will be referred from here on as SC_TET_PT. The highest value obtained when running the adjusted Wallace test was 0.664 (confidence interval of 0.503-0.824) when comparing the ClonalTracker clonal clusters with SC_TET_PT (see Table S6) as ClonalTracker still provides more resolution (higher number of clusters). After a manual screening, three discordant cases were spotted in which ClonalTracker classified isolates into one group, while SC_TET_PT assigned them into different groups based on the transposon type (highlighted in red in Table S5). After re-running the newest TETyper version (the method used in the original publication for the transposon typing step) on these genomes I could confirm that the isolates had the same transposon, and indeed belonged to the same group. Thus, I manually corrected the SC_TET_PT classification and re-run the adjusted Wallace coefficient test (see Table S7), which increased to 0.996 (confidence interval of 0.993-1). This indicates that the plasmid typing step overall does not provide further information as it might already be (at least partially) tackled by ClonalTracker’s whole genome similarity assessment.

**Fig 3.**
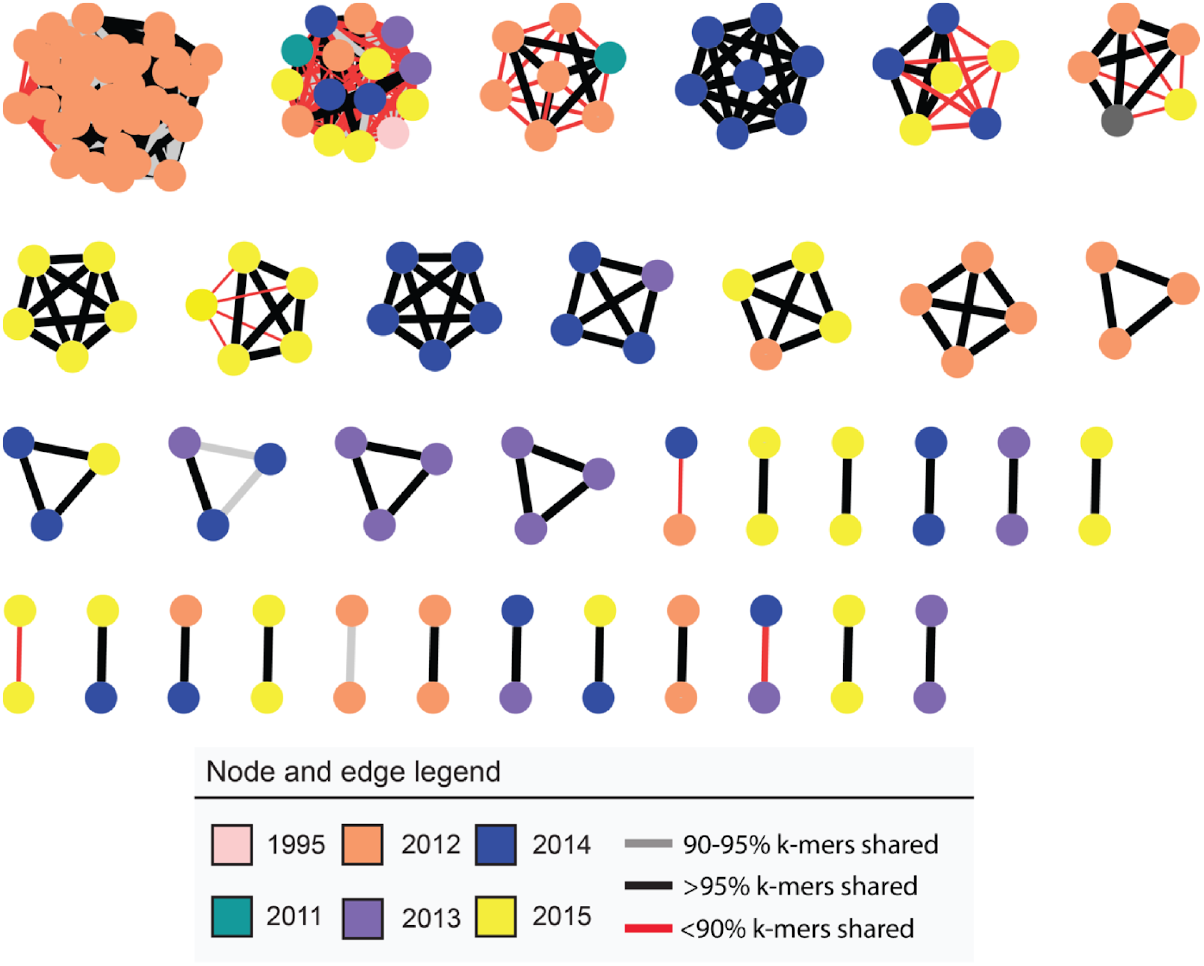
*vanA* dataset clustering based on ClonalTracker results. Genomes (nodes color-coded based on the year of isolation) which share the same transposon are connected by an edge, color-coded based on the percentage of Mash k-mers shared as shown in the legend. Singletons are not included.

In the *vanB*-VRE dataset, ClonalTracker identified a total of 21 different Tn1549-like transposon types. The 5 most frequent types are shared by 23 isolates (see Figure 4, Table S8) while 16 are singletons (unique transposons). In contrast, the original publication only found four different transposon types (TT) using WebACT^29^. All the isolates sharing the same transposon type based on ClonalTracker were also classified under the same TT in the original publication (see Figure 4). However, ClonalTracker was able to find more differences within transposon types and is therefore able to subclassify transposons. For instance, ClonalTracker classified TT2 into two different subgroups (see subnetworks in Figure 4). From the 53 links found by ClonalTracker, 45 were classified as to be clonally-related (represented by black edges in Figure 4). Interestingly, from patient A4 few isolates were sequenced from different body sites (rectum and bile) and based on cgMLST^30^ they belong to different Cluster Types (CT). Isolate A4-2 (CT103) shared the same transposon with the isolates sequenced in April but not with the other isolates sequenced from the same patient (A4-3 and A4-4 which belong to CT24). In line with these findings, ClonalTracker reported that A4-2 does not share the same genomic background as the other isolates in the purple subnetwork (grey and red edges in the subnetwork 1 in Figure 4). This suggests that the isolate A4-2 acquired the *van* resistance from a different source than A4-3 and A4-4.

**Fig 4.**
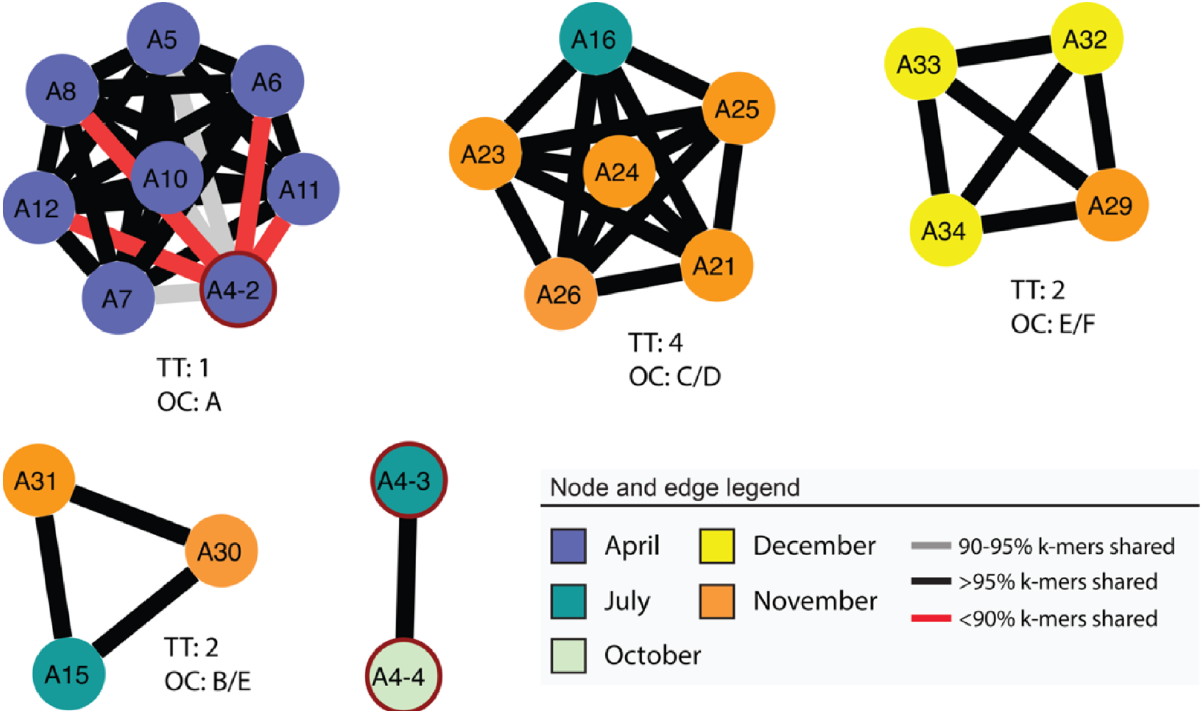
UMCG *vanB* isolates classification based on ClonalTracker. Genomes (nodes color-coded based on the month of isolation) which share the same transposon are connected by an edge, color-coded based on the percentage of Mash k-mers shared as shown in the legend. Singletons are not included. For each subnetwork the original transposon type (TT) classification and outbreak cluster (OC) as described by Zhou et al.^16^ are indicated.

### The *vanB* resistance spreads more frequently via clonal expansion

To better understand the dissemination modes and dynamics of the *vanB* resistance, a collection composed of 323 isolates collected between 2009-2015 from 32 different Dutch hospitals (Figure S1a, Table S9) was analyzed. The number of sequenced genomes was not equally distributed along the years, with 2009-2011 having significantly less sequenced isolates compared to 2012–2015 (Figure S1b). Next, the population structure of this dataset was assessed using MLST, hierBAPS and PopPUNK (see Suppl. results and Figure S1c). The results are also available for exploration at https://microreact.org/project/wRHruR7xfbR24x3bfH3ffY-the-dutch-vanb-dataset.

Subsequently, the dissemination modes of the *vanB* operon within this dataset were assessed by running ClonalTracker. From the 323 isolates, 9 isolates were excluded for either not having the *van* operon (n=3) or harboring both a *vanA* and *vanB* operon (n=6. Table S9). The remaining 314 isolates were analyzed by ClonalTracker, which ran 49,141 pairwise comparisons in total. ClonalTracker identified 125 transposon variants, 34 of which were found in at least two isolates and 91 represented unique transposon types. 223 isolates (shown as nodes in Figure 5) were linked through 2,159 edges (whenever two isolates share the same transposon, see Table S10). Thus each subnetwork can be regarded as a different transposon type. Further results on transposon architecture and diversity can be found in Suppl. material. The *van* typing, the transposon typing and the whole genome similarity analysis showed that most of the linked isolates are clonally related (identical transposon and >95% k-mers shared). This represents 79,3% of the total dissemination events (1,713/2,159 see black edges in Figure 5ab). On the other hand, isolates harboring the same transposon but with different genomic background (<90% k-mers shared), thus suggesting potential horizontal dissemination, only represent 15,6% of the potential dissemination events (336/2159, see red edges in Figure 5ab). The remaining cases (5,1%) were classified as unclear (isolates sharing the same transposon whose genomes share between 90-95% of the k-mers, 110/2159, see gray edges in Figure 5ab). Finally, none of the *vanB* transposons were inserted in a plasmid as confirmed by gplas^31^.

**Fig 5.**
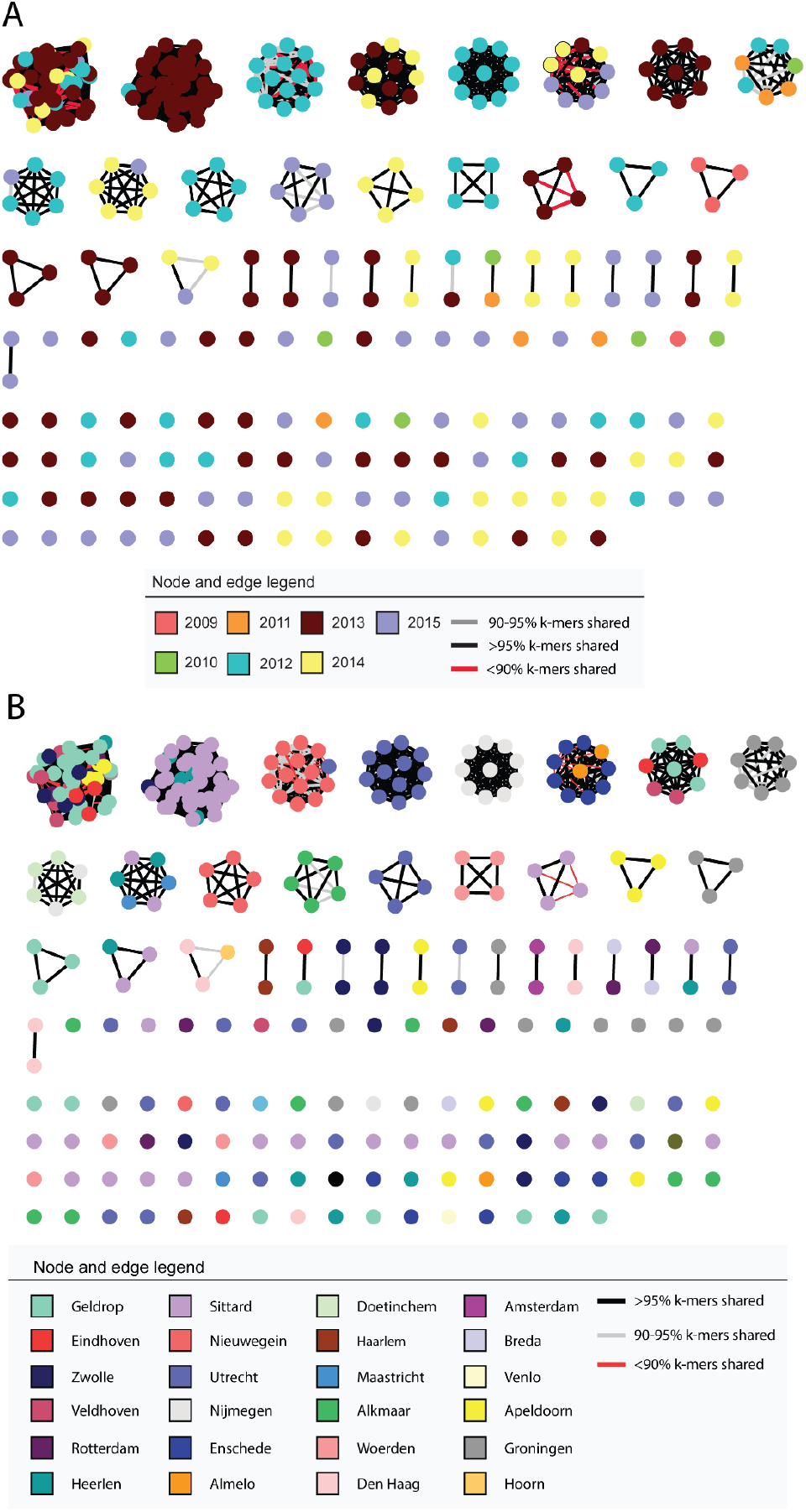
*vanB* dataset clustering based on ClonalTracker results. Genomes (nodes) sharing the same transposon are connected by an edge and are color-coded based on the percentage of Mash k-mers shared as indicated in the legend. a) Nodes are color-coded based on the year of isolation and b) Nodes are color-coded based on the region/city of isolation.

## Discussion

The spread of resistant bacteria especially in healthcare settings may cause increased mortality and morbidity and is also associated with increased healthcare costs^32^. One of the most cost-effective measures to minimize the propagation of bacteria is to do targeted screening and epidemiological typing^33^. With the implementation of WGS in hospitals it is nowadays possible to also track isolates and their mobile genetic elements in a timely manner; a key fact to assess the relatedness between patient’s isolates and proof of nosocomial transmission. As shown in this study, to establish links between VRE isolates and infer that patients are part of the same transmission chain, it is crucial to consider as many layers of genomic information as possible. These different layers of information include clustering based on the core genome alone or in combination with the accessory genomic distance, *van* type and transposon type. Current methods to assess dissemination modes of VRE only focus on one (or few) types of information. In this study, I describe the development of ClonalTracker, which allows to assess the relatedness of any two given VRE genomes. Different datasets have been used to validate ClonalTracker but also to analyze the dissemination modes of an unprocessed *vanB* dataset. In this study multiple *vanB* acquisition events were observed as different transposons were identified among genomically diverse isolates. However, the most frequent dissemination mode is clonal expansion of successful strains. At the regional and hospital level more complex dissemination patterns are also seen supporting other studies which evinced *vanB* horizontal dissemination across different hospitals^34^,^35^.

ClonalTracker uses information from several complementary methods to assess clonality. Specifically, to assess the *van* transposon, this tool uses a reference sequence dependent method. To systematically identify *vanA* and *vanB*-harboring transposons a reference including the minimum number of genes common in all *van* transposons is used: the *van* operon plus excisase and integrase. A mapping step with the reference transposon is performed to determine the transposon start and end coordinates. However, this implies that (in some cases) the transposon boundaries might not be accurately defined as genes flanking the matching region are ignored. Moreover, using a reference-based approach could hinder the identification of novel and unrelated transposon types. Although reference-free approaches would be ideal to not constrain the transposon identification, using a rather “simple” transposon architecture still allows to accurately identify transposons which differ at the SNP and gene content level. Additionally, ClonalTraker has been proven to be more fine-grained than others (or combinations of them) such as TETyper, gplas, Mash and WebACT; and helped elucidate a large *vanB* transposon collection.

Assessing the sequence identity of all the mobile genetic elements involved in the vancomycin resistant phenotype is important to evaluate the relatedness between VRE isolates. However, ClonalTracker does not specifically take into account the role of the plasmids in VRE dissemination. As shown in this study, this limitation is alleviated by whole genome comparison, where the whole genome sequence (including plasmids) is randomly subsampled and compared. It is important to highlight that ClonalTracker outputs all intermediate and final results, thus allow users to perform follow-up analyses as for instance assess the transposition site and flanking regions of the transposon to further confirm (or not) clonal relatedness, visualize transposon sequence similarity in a different way and/or check the presence of plasmids.

Networks have been shown to be very useful in epidemiology to trace back transmission routes using contact networks^36^, visualize epidemic spreading^37^, assess pathways of gene transmission and the antibiotic resistance genes shared between organisms^38^, for instance. Here, I used networks to visualize the relationships between isolates, transposon diversity, transposon type abundance and isolate genome similarity, as well as isolate metadata. This allows to uncover links which otherwise (for e.g., using phylogenies) would have been very difficult to grasp. Networks can be very informative as there are many different ways to annotate them and highlight specific features of each node and edge. Nonetheless, interpretation of these links might sometimes be difficult. For instance, edge directionality is hard to establish specially in the absence of extensive metadata and might be key to determine the origin of a given outbreak retrospectively. Another limitation encountered when using networks based on transposon identity is that edges are not (always) equivalent to transmission events. When clone A transfers a given transposon to clone B and subsequently spreads the transposon clonally to B1, B2 and B3, in our network, clone A would still be linked to B1, B2 and B3 despite there should not be a direct edge between them if only interested in transmission events. Thus, to confirm between clonal and horizontal gene transmission within our dataset it is required more detailed metadata regarding isolations dates, patients ward, among others, and expert assessment of the networks.

In summary, I have designed a method to elucidate links between VREs using the assemblies obtained from short-read sequencing as input: the preferred sequencing technology for such studies to date^39^. Nonetheless, this technology also comes with its limitations. For instance, full reconstruction of mobile genetic elements can be challenging given the number of short repeated sequences but also to determine the genetic context where the resistance genes are located can be difficult. Thus, as ClonalTracker takes the already assembled genome as input, it is important that it is as complete as possible. If the transposon is split into multiple contigs, ClonalTracker scaffolds them whenever they can still be captured. However, rearrangements, deletions or insertions that are not present in the reference transposon used would be harder to include. Therefore, the more accurately reconstructed the transposon is (but also the whole genome) the more accurate results ClonalTracker will provide. To ensure it, a good solution is to use hybrid assemblies obtained from combining both short and long-sequencing reads. This has proven to be a successful method to not only recover mobile genetic elements^40^ but also to obtain more accurate and complete genomes^41^. Overall, the algorithm has been designed to assess clonal relatedness between VRE isolates. However, creating new databases and choosing the corresponding reference sequences the tool could also be implemented to get further insights into dissemination modes of other antimicrobial resistances and bacteria.

## Supporting information

Supplementary Results

## Acknowledgments

The author thanks the guidance provided by Janetta Top, Rob Willems and Anita Schürch throughout the duration of the project. This project has been funded by ZonMw.

## Data availability

The raw reads of the VRE isolates used in this article are available at the following ENA accession numbers: PRJEB28495 and PRJEB25590. The metadata of the former can also be accessed from here: https://microreact.org/project/BJKGTJPTQ.

## Code availability

The ClonalTracker command line version can be found here: https://github.com/victoriapascal/clonaltracker

