## Supplementary Results for "ClonalTracker: a tool to elucidate dissemination patterns between vancomycin-resistant *Enterococcus faecium* isolates"

### Population structure of the *vanB* dataset

The dataset composed of 323 *vanB* isolates includes isolates from different geographic locations, isolated in different years but also genomically diverse (see Figure S1). All different typing/clustering tools revealed a very diverse genomic background among the isolates. Based on hierBAPS clustering, 321 isolates (99.38%) were found in the hospital-associated clade A1 and only two isolates (E7838 and E8029) were not part of the clade A1 (see Figure 4c). Multi-Locus Sequence Type (MLST) classified the 323 *vanB* isolates into 12 different groups with the majority of the isolates belonging to either Sequence Type (ST) 117 (n=194) and ST192 (n=71). To further classify them using core and accessory gene distances PopPUNK was run, which allowed a more fine-grained classification and grouped the 323 isolates into 109 different clusters. Most of the isolates were classified in either cluster 1 (n=61) or 2 (n=44) (Figure S1c).

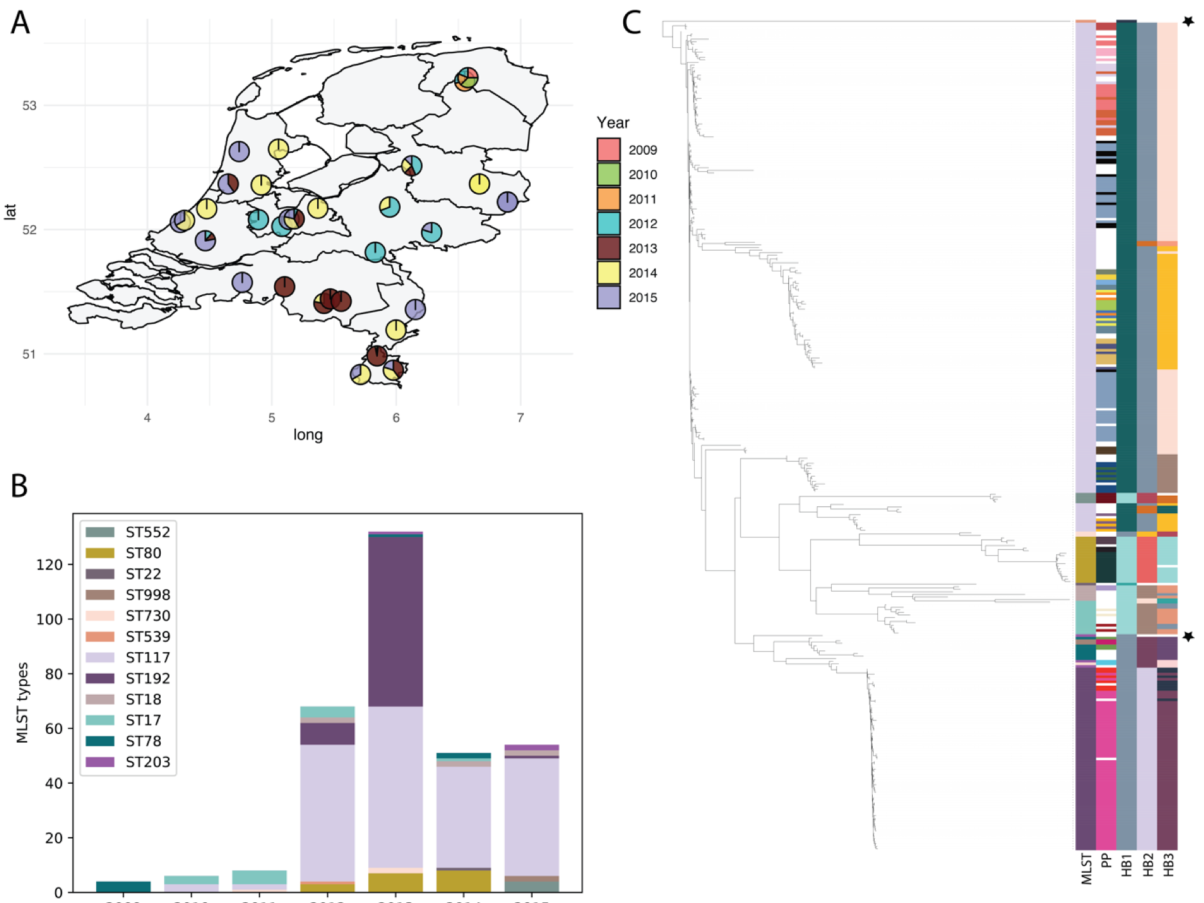


**Fig S1. Distribution and population structure of the *vanB* dataset.** a) Relative number of isolates sequenced per year and hospital b) Stacked bar plot showing the year of isolation and MLST of the *vanB*-VRE isolates. c) Core genome-based tree generated by PopPUNK. The tree was annotated with iTOL to include five panels on the right which represent the MLST type, the PopPUNK clustering (PP) using core and accessory gene distances (only clusters of size > 1 are shown) and the hierBAPS1, -2 and -3 respectively. Stars indicate the two clade A isolates (E7838 and E8029).

**Transposon architecture and diversity within the *vanB* dataset**

ClonalTracker identified a total of 125 transposon variants, of which 34 were found in at least two isolates and 91 represented unique transposon types (singletons). This indicates that the *vanB* resistance has been introduced or has evolved multiple times along the course of this study. Variation between transposon-types mostly includes differences at SNP level (see Figure S2), but there are also transposon types which have different architectures (see branch distances in Figure S2a) indicating that some transposon types are more distantly related than others. By selecting a few of them (see Figure S2b) it evinces that the transposons differ in sequence length but also in gene's sequence identity. More specifically, the ones which carry most mutations are vanS (sensor), vanY (carboxypeptidase), vanH (dehydrogenase), vanB (ligase) and vanX (dipeptidase).

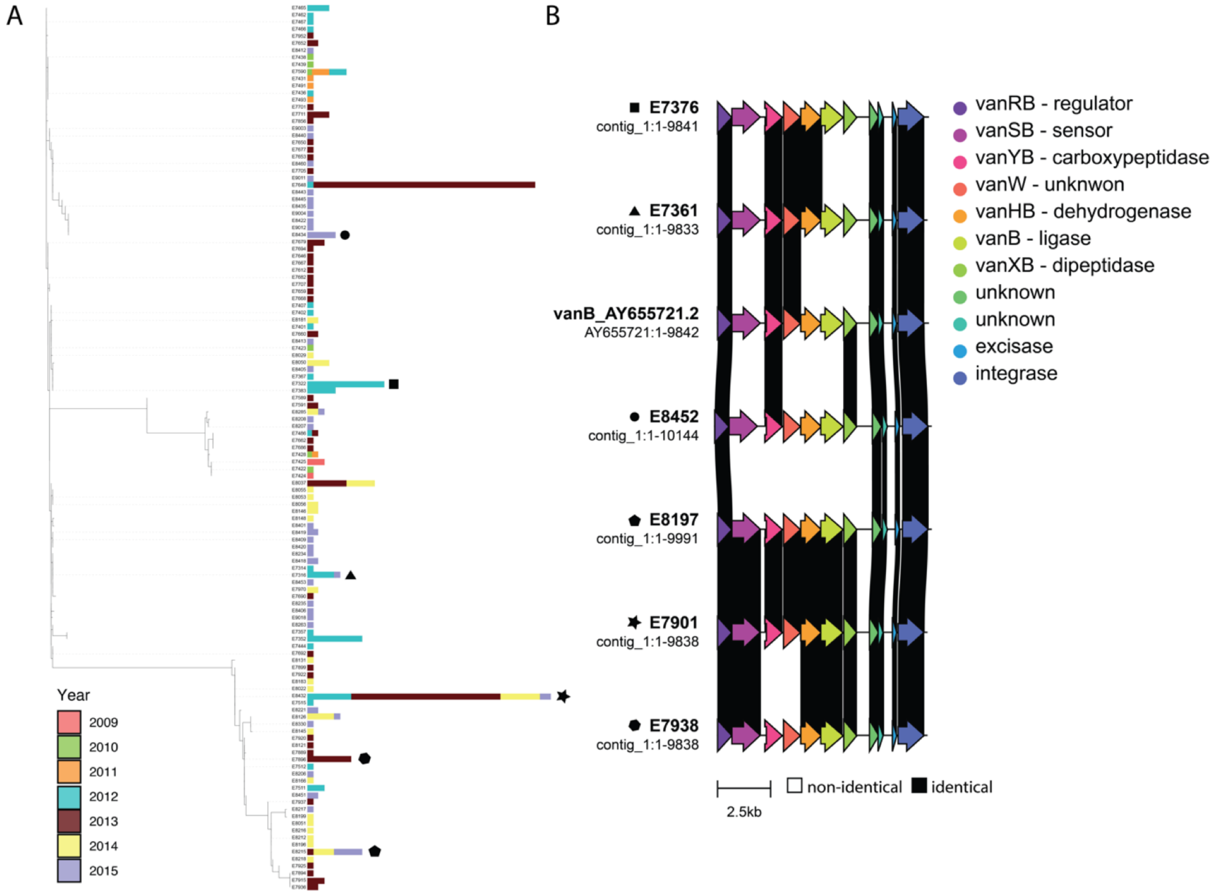


**Fig S2: *vanB* transposon diversity**. a) from each transposon type a random sequence was chosen as representative for a multiple sequence alignment with MAFFT[1] and subsequently build a phylogeny using FastTree[2]. b) Six random representatives were chosen from the tree. The annotated sequences retrieved by ClonalTracker including the ClonalTracker *vanB* reference sequence (*vanB* AY655721.2) were used as input for clinker. Each transposon type selected in panel b is identified with the same shape in both panel a and b.
